# Cell type specific IL-27p28 (IL-30) deletion uncovers an unexpected regulatory function of IL-30 in autoimmune inflammation

**DOI:** 10.1101/2022.01.03.474823

**Authors:** Dongkyun Kim, Sohee Kim, Myung-su Kang, Zhinan Yin, Booki Min

## Abstract

IL-27 is an IL-12 family cytokine with immune regulatory properties, capable of modulating inflammatory responses, including autoimmunity. While extensive studies investigated the major target cells of IL-27 mediating its functions, the source of IL-27 especially during tissue specific autoimmune inflammation has not formally been examined. IL-27p28 subunit, also known as IL-30, was initially discovered as an IL-27-specific subunit, and it has thus been deemed as a surrogate marker to denote IL-27 expression. However, IL-30 can be secreted independently of Ebi3, a subunit that forms bioactive IL-27 with IL-30. Moreover, IL-30 itself may act as a negative regulator antagonizing IL-27. In this study, we exploited various cell type specific IL-30-deficient mouse models and examined the source of IL-30 in T cell mediated autoimmune neuroinflammation. We found that IL-30 expressed by infiltrating and CNS resident APC subsets, infiltrating myeloid cells, microglia, and astrocytes, is key limiting the inflammation. However, dendritic cell-derived IL-30 was dispensable for the disease development. Unexpectedly, in cell type specific IL-30 deficient mice that develop severe EAE, IL-30 expression in the remaining wild-type APC subsets is disproportionately increased, suggesting that increased endogenous IL-30 production may be involved in the severe pathogenesis. In support, systemic recombinant IL-30 administration induced severe EAE. Our results demonstrate that dysregulated endogenous IL-30 expression may interfere with immune regulatory functions of IL-27, promoting encephalitogenic inflammation in vivo.

## Introduction

IL-27 is an IL-12 family heterodimeric cytokine composed of p28 (also known as IL-30) and Ebi3 subunits, and binds the IL-27 specific receptor, a heterodimeric surface receptor complex made of IL-27Rα and gp130^1,2^ IL-27 mediates highly diverse, even opposing, pro- and anti-inflammatory roles by supporting Tbet/IFNγ expression in developing Th1 cells and by inhibiting Rorc expression and Th17 differentiation, respectively^3–5^. Another well appreciated anti-inflammatory function of IL-27 operates through IL-10 induction from activated CD4 T cells^6,7^. IL-10-producing Foxp3^-^ helper T (known as Tr1) cells are thought to play an important role in suppressing inflammation and maintaining tolerance in many conditions^8,9^. Indeed, mice deficient in IL-27Rα subunit are highly susceptible to Th17-mediated autoimmune inflammation, experimental autoimmune encephalomyelitis (EAE), and the susceptibility is thought to be attributed to the deficit in the development of IL-10-producing CD4 T cells^10^. However, IL-27 directly acting on Foxp3^+^ Treg cells supports Treg cells’ ability to suppress autoimmune inflammation via Lag3-dependent and Tr1-independent mechanisms^11^.

Being the IL-27-specific subunit, IL-30 was measured as a surrogate to assess IL-27 production. The primary source of IL-30 is cells of myeloid origin, including monocytes, macrophages, and dendritic cells^2^ Signals triggering IL-30 secretion are mostly of innate immunity. TLR3 and TLR4 have previously been shown to trigger IL-30 expression in dendritic cells via an IRF3-dependent mechanism^12^, and IFNγ signal can augment the expression^13,14^. IFNβ, a widely used immune suppressive cytokine for the treatment of autoimmunity, is another cytokine inducing IL-30 production and inhibiting Th17 differentiation^15^. However, little is known regarding the precise source and immune regulatory functions of IL-30 especially during tissue specific autoimmune inflammation, such as EAE.

There is emerging evidence that IL-30 may exhibit an immune regulatory function distinct from that of IL-27^16,17^ Caspi and colleagues utilized IL-30-overexpressing mouse model to show that IL-30 inhibits T cell differentiation to Th1 and Th17 lineage cells and the development of autoimmunity^18^. IL-30 also negatively regulates humoral and cellular responses during parasite infection, independent of its role as an IL-27 subunit^19^. In the context of tumorigenesis, IL-30 has been suggested to play a pro-tumorigenic role, in part by supporting cancer stem-like cell survival, vascularization, and proliferation^20,21^. However, the nature of such pro-inflammatory and pro-tumorigenic functions of IL-30 remains largely unclear^22^.

The current study aimed at identifying the source of IL-30 and its potential role during autoimmune inflammation in the central nervous system (CNS). We utilized various cell type specific *Il27p28*^-/-^ mouse models and found that IL-30 expressed by myeloid cells, microglia, and astrocytes but not by dendritic cells plays an important role in limiting autoimmune inflammation. Unexpectedly, we noted that exacerbated inflammation seen in those cell type specific IL-30-deficient mice was associated with drastic overexpression of *Il27p28* mRNA in otherwise unaffected wild type CNS-infiltrating and resident APC subsets, possibly resulting in disproportionate IL-30 production. Systemic administration of recombinant IL-30 alone into mice with ongoing EAE similarly aggravated the disease progression, suggesting that IL-30 itself can enhance encephalitogenic immune activity. Increased IL-30 expression was associated with decrease in Treg cell expression of Lag3, an indicative of IL-27 signals, suggesting that IL-30 may antagonize IL-27’s action on Treg cells in vivo. Yet, IL-30 showed no measurable biological activity on activated T cells as determined by its ability to phosphorylate Stat1 and Stat3 or to antagonize IL-27 activity in vitro. Taken together, IL-30 may exert a distinct function to antagonize IL-27 and/or to support autoimmune inflammatory T cell responses in vivo.

## Results

### Cytokine mRNA expression in EAE

We first measured cytokine gene expression in the brain and spinal cords during EAE. Three time points were chosen: days 8, 14, and 21 post induction, representing the disease onset, peak, and partial recovery, respectively. *Il27p28* mRNA expression mirrored the disease activity, and it peaked at day 14, showing >10-fold increase in both tissue sites compared to those of naïve mice (Fig 1a). The *Ebi3* subunit mRNA expression displayed a similar pattern as the *Il27p28* (Fig 1a). *Il12p40* mRNA expression in the spinal cord was markedly increased with the similar kinetics, reaching ~100-fold increase over naïve tissue, although its expression in the brain was not observed (Fig 1a). On the other hand, *Il12p35* or *Il23p19* mRNA expression only slightly increased (data not shown). Expression of inflammatory cytokines, such as, *Tnfa, Ifng, Il17a, Il1b*, and *Il6*, as well as of key transcription factors, *Tbx21* and *Rorc*, followed the similar pattern (Fig 1b and 1c). Foxp3 mRNA expression substantially increased at the peak of the disease and was maintained thereafter, demonstrating Treg accumulation in the tissue, likely involved in inflammatory resolution^23^. We also measured inflammatory chemokines, such as *Ccl2, Ccl3, Ccl7, Cxcl1, Cxcl9*, and *Cxcl10*. Again, the expression pattern showed similar kinetics, with the greater magnitude in the spinal cord (Supp Fig 1). Myeloid cells capable of presenting antigens, including macrophages and dendritic cells, are the primary source of IL-12 family cytokines including IL-27. To examine relative sources of each cytokine during autoimmune inflammation in the CNS, we FACS sorted different APC subsets from the inflamed CNS tissues; CD45^high^ CD11b^high^ infiltrating myeloid cells, CD45^int^ CD11b^high^ microglia, and CD45^low^ cells that include astrocytes and oligodendrocytes, at the peak of the disease, and cytokine gene expression was determined. While both infiltrating myeloid cells and microglia similarly expressed all the tested IL-12 family cytokines, the level of *Il27p28* and *Ebi3* mRNA expression was particularly greater than any other subunits examined (Fig 1d). These results prompted us to investigate the central source of IL-27, especially IL-27p28 (referred to as IL-30 hereafter) subunit during the development of autoimmune neuroinflammation.

**Figure 1.**
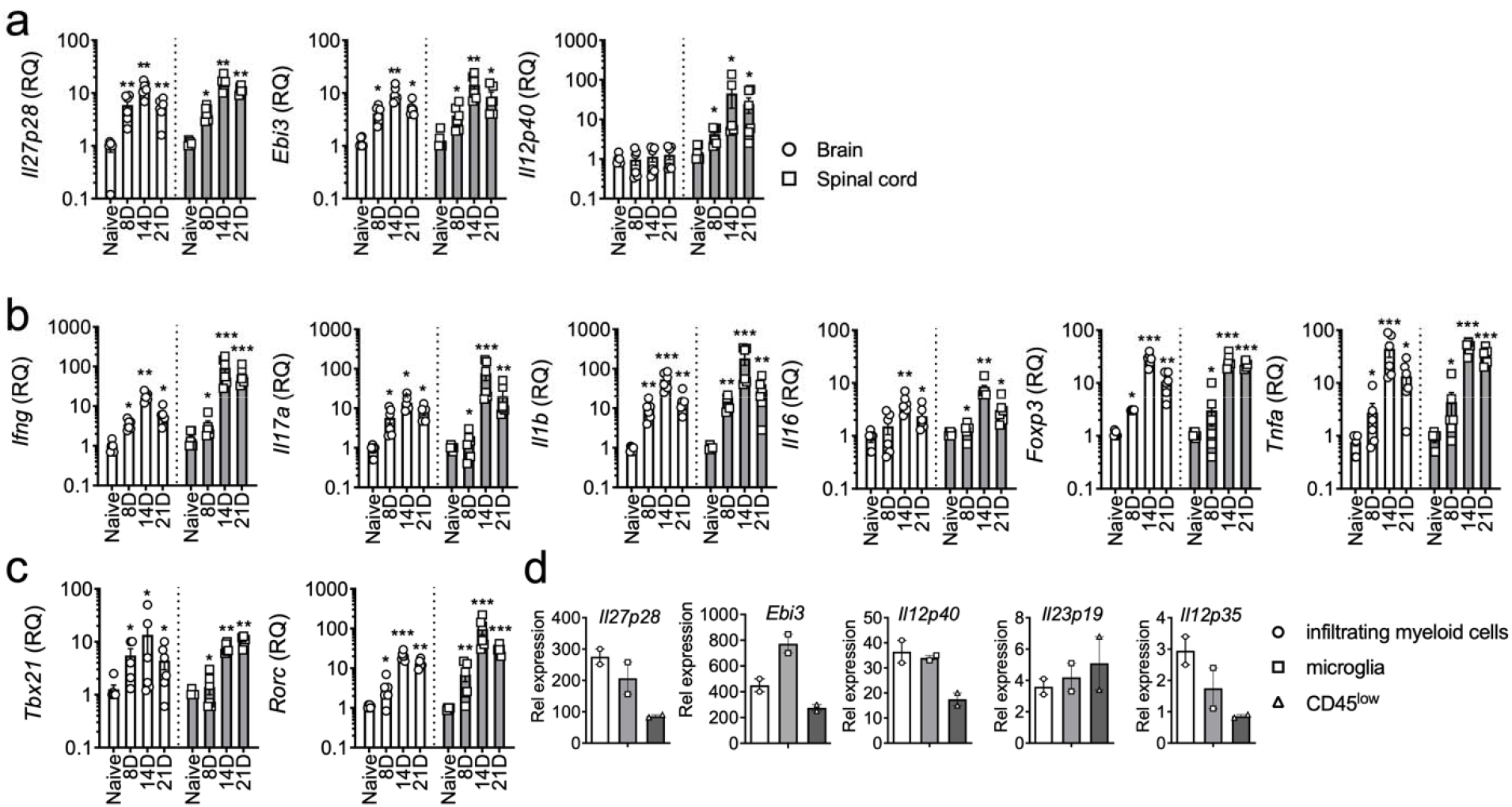
Cytokine gene expression in the CNS during EAE. (a-c) EAE was induced in C57BL/6 mice as described in the Methods. RNA was isolated from the brain and spinal cords at disease on set (day 8 post immunization), acute phase (day 14 post immunization) and remission phase (day 21 post immunization). n = 4-6 per group. mRNA expression of IL-12 family genes, cytokines and transcription factors were measured by qRT-PCR. Data were normalized by *Gapdh* gene expression and compared to that of naive mice. (d) CD45^high^ CD11b^high^ (infiltrating myeloid cells), CD45^int^ CD11b^high^ (microglia) and CD45^low^ (including astrocyte and oligodendrocyte) cells sorted from the CNS at the peak of disease (day 17 post immunization) and expression of the indicated genes was measured by qPCR. Gene expression was normalized by *Gapdh* and compared to that of naïve mice. The results shown represent two independent experiments. *p < 0.05; **p < 0.01; ***p < 0.001; as determined by Mann-Whitney nonparametric test.

### Myeloid cell-derived IL-30 regulates encephalitogenic immune responses

Cells of myeloid origin, especially monocytes and macrophages, are an important source of IL-30^1,24^. To interrogate the contribution of myeloid cell-derived IL-27 to EAE pathogenesis, we utilized myeloid cell-specific IL-30^-/-^ (LysM^Cre^ *Il27p28*^fl/fl^) mice. The lack of *Il27p28* mRNA expression in macrophages in LysM^Cre^ *Il27p28*^fl/fl^ mice was validated by qPCR (Supp Fig 2). Myeloid cell-specific IL-30^-/-^ mice developed severe EAE, although they appear to recover from the initial paralysis similarly to that of wild type mice (Fig 2a). Severe acute disease was further reflected by increased CD4 T cells infiltrating the CNS at the peak of the disease (Fig 2b). CNS accumulation of Foxp3^+^ Treg cells was significantly greater in these mice, although Treg cell proportion or Foxp3 expression was found comparable (Fig 2c). Treg cell expression of ICOS, GITR, and CD25 remained unchanged regardless of myeloid cell-derived IL-30 (data not shown). CD4 T cells expressing inflammatory cytokines were substantially increased in the CNS tissues of LysM^Cre^ *Il27p28*^fl/fl^ mice (Fig 2d), consistent with severe EAE phenotypes in these mice. In support, serum cytokine levels, especially IL-6, IFNγ, IL-17A, and TNFα, were significantly increased in myeloid cell specific IL-30^-/-^ mice (Fig 2e). mRNAs for encephalitogenic cytokines and key transcription factors were similarly elevated in these mice (Fig 2f), and inflammatory chemokine expression in the CNS tissues was similarly elevated in mice with myeloid cell specific IL-30 deletion (Supp Fig 3). CNS expression of *Il27p28* and *Il12p40* mRNAs was greater in the absence of myeloid cell-derived IL-30, which may be attributed to elevated expression of IL-30-inducing cytokines such as IFNγ (Fig 2f). *Il23p19* and *Il12p35* mRNA expression was similar between the groups (Fig 2f). Increased *Il27p28* mRNA expression in LysM^Cre^ *Il27p28*^fl/fl^ mice was further examined by comparing the expression in FACS sorted CNS APC subsets. Unexpectedly, we observed that *Il27p28* mRNA expression was drastically increased in microglia and CD45^low^ cells in this condition (Fig 2g). *Il27p28* mRNA expression in infiltrating myeloid cells was not found, corroborating IL-30 deficiency of myeloid cells (Fig 2g). *Ebi3* mRNA expression was also increased in microglia and CD45^low^ cells (Fig 2g), while myeloid cell expression of the *Ebi3* mRNA remained unchanged. Therefore, myeloid lineage cells could be an important source of IL-27 (and IL-30), capable of modulating EAE pathogenesis.

**Figure 2.**
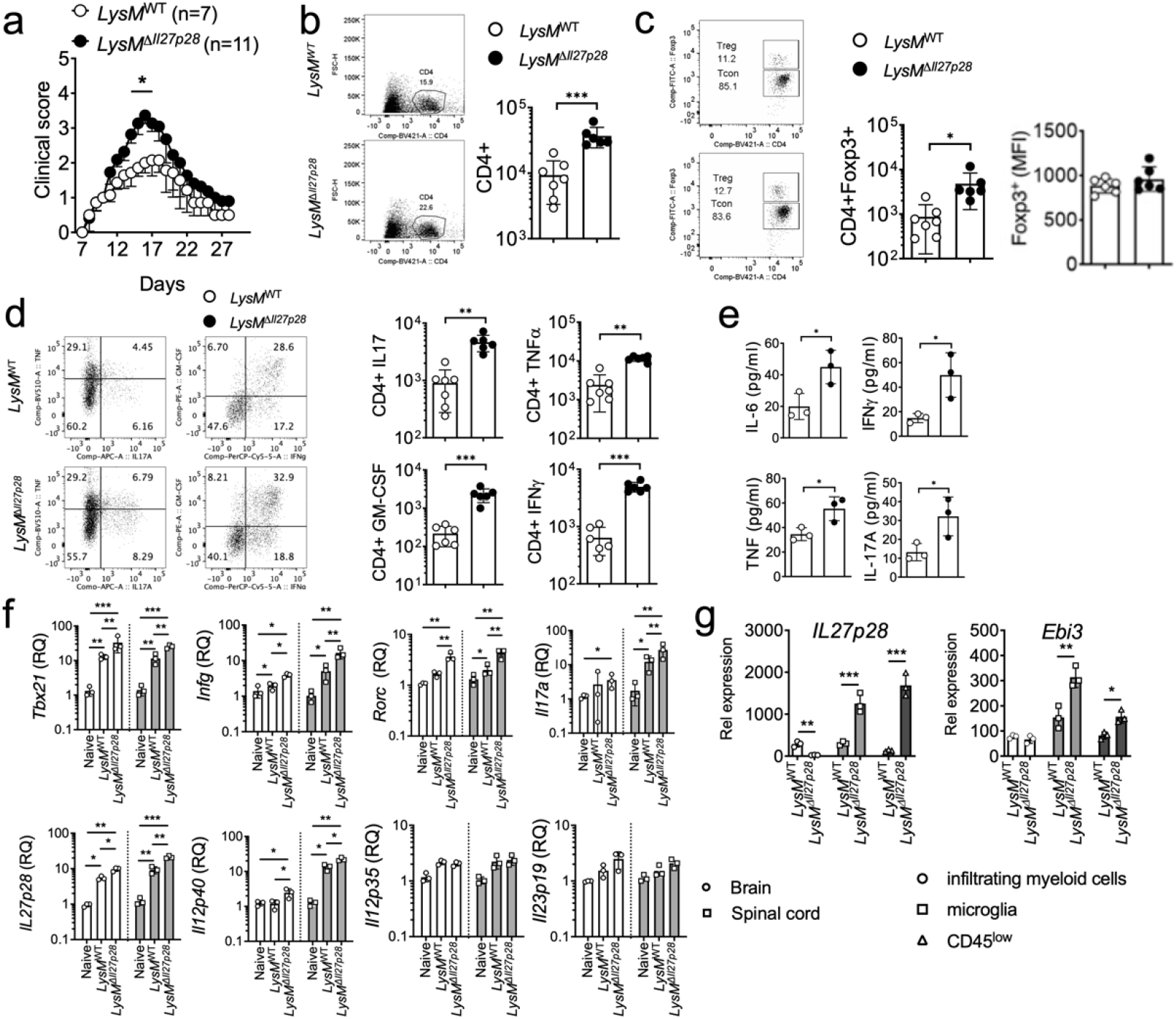
EAE in myeloid cell-specific *Il27p28*^-/-^ mice. LysM^*WT*^ (n = 7) and LysM^Cre^ *Il27p28*^fl/fl^ (LysM^Δ*Il27p28*^) (n = 11) mice were induced for EAE. (a) EAE clinical score. (b-c) The numbers of CNS infiltrating CD4^+^ and CD4^+^Foxp3^+^ Treg cells, and the mean fluorescence intensity (MFI) of Foxp3 was determined by flow cytometry at day 17 post immunization. (d) Flow cytometry analysis of GM-CSF, IFNg, IL-17, and TNFa CD4^+^ **T** cells from the CNS of EAE mice (day 17 post immunization). (e) qPCR analysis of the indicated mRNAs in the brain and spinal cords from naïve, LysM^WT^, and LysM^Δ*Il27p28*^ mice 17 days post immunization. Gene expression was normalized by *Gapdh* and compared to that of naïve mice. n = 3 per group. (f) The levels of IL-6, IFNγ, TNFα and IL-17A in the serum of EAE mice (day 17 post immunization) were measured using Cytometric Bead Array. Each serum sample was analyzed in duplicates. (g) qPCR analysis of the indicated mRNAs in freshly sorted CD45^high^ CD11b^high^ (infiltrating myeloid cells), CD45^int^ CD11b^high^ (microglia) and CD45^low^ (astrocyte an**d** oligodendrocyte) cells from LysM^WT^ or *LysM*^Δ*Il27p28*^ mice 17 days post immunization. n = 3 per group. *p < 0.05; **p < 0.01; ***p < 0.001; as determined by Mann-Whitney nonparametric test.

### DC-derived IL-30 is dispensable

IL-27 produced by DCs promotes the generation of IL-10^+^ CD4 T cells capable of attenuating autoimmune inflammation^25^. Utilizing the IL-30-floxed mouse model, it was concluded that DC-derived IL-27 plays a role in antitumor immunity by regulating NK and NKT cell recruitment and activation^26,27^. To test the role of DC-derived IL-27 in EAE, DC-specific IL-30^-/-^ (CD11c^Cre^ *Il27p28*^fl/fl^ mice were used. qPCR analysis validated DC-specific IL-30 deficiency in these mice (Supp. Fig 2). Despite the involvement of DC-derived IL-27 in T cell immunity, we found that DC-specific IL-30 deletion did not affect EAE pathogenesis, as both disease onset and the clinical severity remained comparable to those of wild type control mice (Fig 3a). CD4 T cells infiltrating the CNS tissues were similar in both proportions and absolute numbers (Fig 3b). Likewise, Foxp3^+^ Treg cell accumulation in the CNS was also comparable (Fig 3c). Treg cell expression of surface markers associated with the suppressive function, namely, ICOS, GITR, and CD25, was similar regardless of DC-derived IL-30 (Fig 3d). We then measured CD4 T cell expression of encephalitogenic cytokines by flow cytometry. As shown in Fig 3e, intracellular expression of GM-CSF, IFNγ, IL-17, and TNFα in CNS infiltrating CD4 T cells was similar in both proportions and absolute numbers. Consistent with these findings, expression of *Ifng, Il17a*, and *Il1b* mRNA in the CNS tissues was also similar between WT and DC-specific IL-30^-/-^ groups (Fig 3f). Likewise, serum cytokine levels measured by cytokine bead array (CBA) were comparable between the groups (Fig 3g). Inflammatory chemokine and IL-12 family cytokine expression in the tissues was similar between the groups (Supp Fig 4a and 4b). Lastly, CNS APC subsets were FACS sorted as above, and IL-12 family cytokine gene expression was determined. Consistent with the EAE severity and overall immune responses, we found no differences in cytokine gene expression between the groups (Fig 3h and not shown). Therefore, DC-derived IL-30 plays little role in regulating encephalitogenic immune responses.

**Figure 3.**
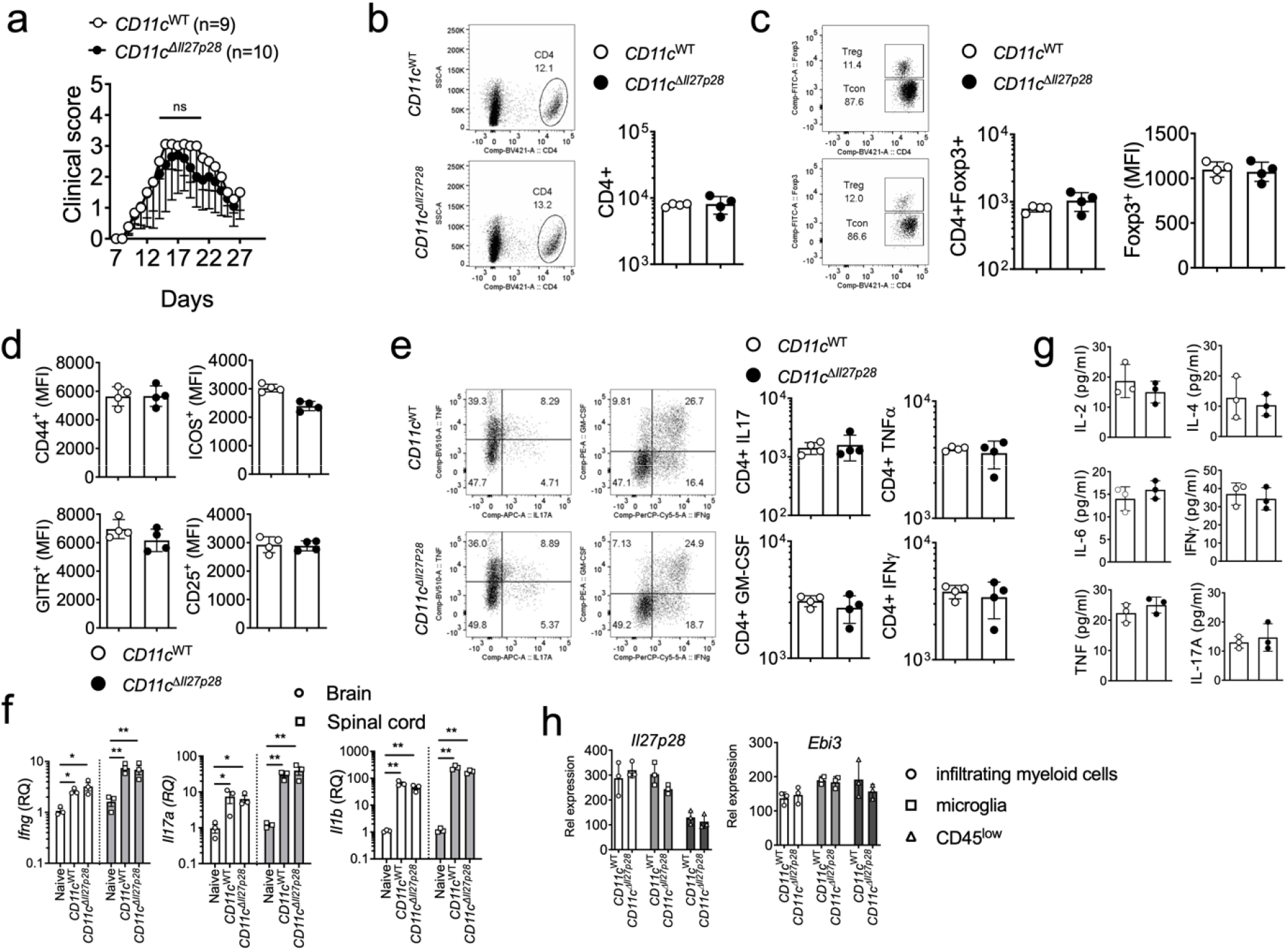
EAE in DC-specific *Il27p28*^-/-^ mice. CD11c^WT^ (n = 9) and CD11c^Cre^ *Il27p28*^fl/fl^ (CD11c^*ΔIl27p28*^) (n = 10) were induced for EAE. (a) Time course of the development of EAE. (b-d) The numbers of CNS-infiltrating CD4^+^ and CD4^+^Foxp3^+^ Treg cells, and the mean fluorescence intensity (MFI) of Foxp3, CD44, ICOS, GITR and CD25 were determined by flow cytometry at day 17 post immunization. (e) Flow cytometry analysis of GM-CSF, IFN-γ, IL-17, and TNFα CD4^+^ T cells from the CNS of EAE mice (day 17 post immunization). (f) RNAs isolated from the brain and spinal cords at day 17 post immunization were analyzed for the expression of *Ifng, Il17a*, and *Il1b*. n = 3 per group. Gene expression was normalized by *Gapdh* and compared to that of naïve mice. (g) The levels of IL-2, IL-4, IL-6, IFNg, TNFa and IL-17A in the serum of EAE mice (day 17 post immunization) were measured using the Cytometric Bead Array. Each serum sample was analyzed in duplicates. (h) qPCR analysis of the indicated mRNAs in freshly sorted CD45^high^ CD11b^high^ (infiltrating myeloid cells), CD45^int^ CD11b^high^ (microglia) and CD45^low^ (astrocyte and oligodendrocyte) cells from CD11c^WT^ and CD11c^*ΔIl27p28*^ mice with EAE. *p < 0.05; **p < 0.01; ***p < 0.001; as determined by Mann-Whitney nonparametric test.

### Microglia-derived IL-30 plays a similar regulatory role in EAE

Microglia are resident CNS glial cells capable of producing IL-27^28,29^ Since LysM^Cre^-mediated gene targeting can occur in microglia as well^30^, we investigated the role of microglia-derived IL-27. To target microglia expression of IL-30, Cx3cr1^Cre^ *Il27p28*^fl/f^ mice were used^31^. Microglia specific IL-30^-/-^ mice exhibited severe EAE analogous to myeloid cell specific IL-30^-/-^ mice (Fig 4a). CNS infiltration of total CD4 and Foxp3^+^ Treg cells was significantly increased in these mice, supporting severe EAE phenotypes (Fig 4b and 4c). Treg cell associated surface marker and Foxp3 expression remained unchanged in microglia-specific IL-30^-/-^ mice (Fig 4c and data not shown). CNS infiltrating CD4 T cell expression of inflammatory cytokines was markedly increased, further supporting greater susceptibility of these mice (Fig 4d). We measured inflammatory cytokine and chemokine gene expression by qPCR and confirmed that microglia-specific IL-30 deficiency results in greater increase of the expression (Fig 4e and data not shown). *Il27p28* mRNA expression in the CNS of microglia-specific IL-30^-/-^ mice was significantly elevated compared to that of wild type mice (data not shown). Also observed was that *Il27p28* mRNA expression in FACS sorted different CNS APC subsets, i.e., infiltrating myeloid cells and CD45^low^ cells, was drastically increased in microglia-specific IL-30^-/-^ mice (Fig 4f). The lack of *Il27p28* mRNA expression in sorted microglia further validated microglia-specific IL-30 deficiency (Fig 4f). Modest increase of *Ebi3* mRNA expression was also seen in infiltrating monocytes and CD45^low^ cells but not in microglia (Fig 4f). Therefore, microglia-derived IL-30 also appears to play a regulatory role in EAE pathogenesis.

**Figure 4.**
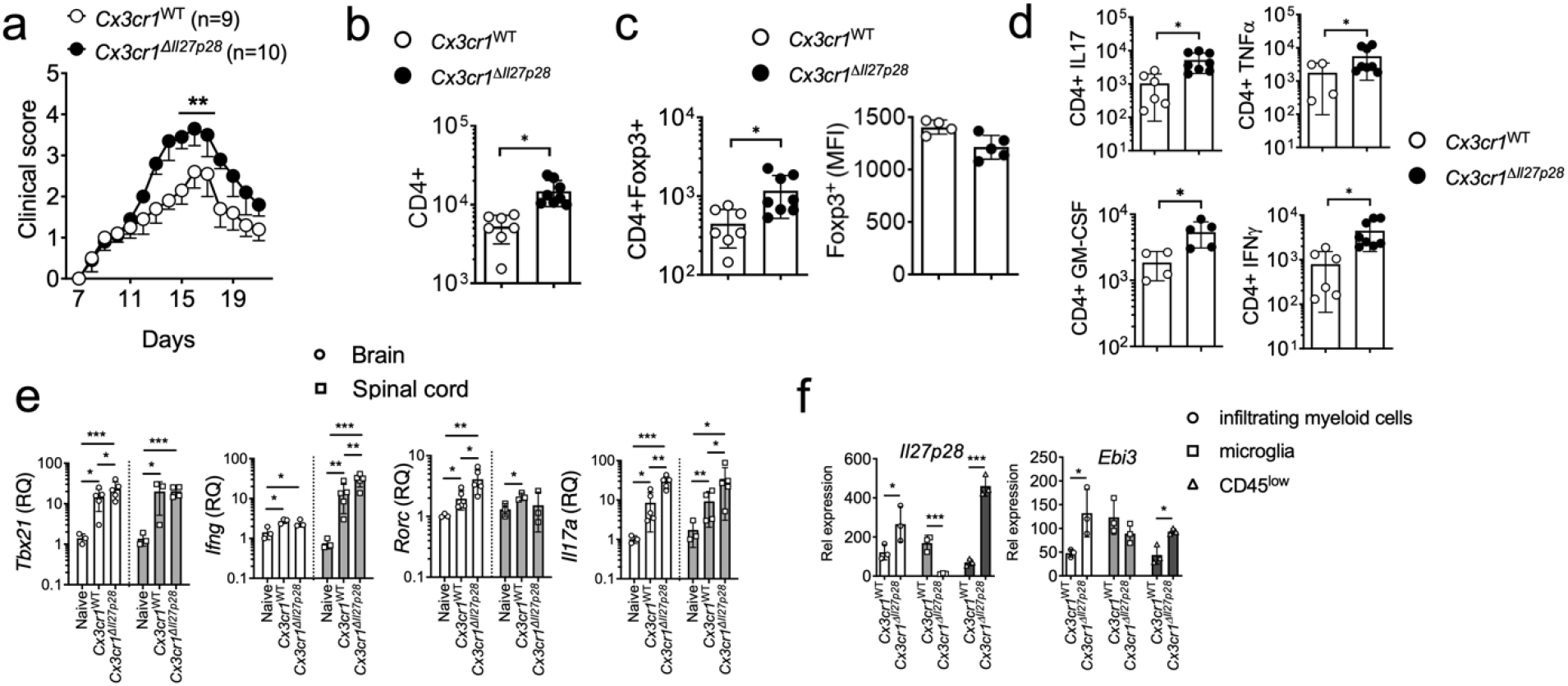
EAE in microglia-specific *Il27p28*^-/-^ mice. Cx3cr1^WT^ (n = 9) and Cx3cr1^Cre^ *Il27p28*^fl/fl^(Cx3cr1^Δ*Il27p28*^) (n = 10) were induced for EAE. (a) EAE clinical scores. (b-c) Total numbers of CNS-infiltrating CD4^+^ and CD4^+^Foxp3^+^ Treg cells, and the mean fluorescence intensity (MFI) of Foxp3 was determined by flow cytometry at day 17 post immunization. (d) Flow cytometry analysis of GM-CSF, IFNγ, IL-17, and TNFα CD4^+^ T cells from the CNS of EAE mice (day 17 post immunization). (e) qPCR analysis of the indicated mRNAs in the brain and spinal cords from naïve, Cx3cr1^WT^, and Cx3cr1^Δ*Il27p28*^ mice 17 days post immunization. Gene expression was normalized by *Gapdh* and compared to that of naïve mice. n = 3-5 per group. (f) qPCR analysis of the indicated mRNAs in freshly sorted CD45^high^ CD11b^high^ (infiltrating myeloid cells), CD45^int^ CD11b^high^ (microglia) and CD45^low^ (including astrocyte and oligodendrocyte) cells from Cx3cr1^WT^ or Cx3cr1^Δ*Il27p28*^ mice 17 days post immunization. n = 3 per group. *p < 0.05; **p < 0.01; ***p < 0.001; as determined by Mann-Whitney nonparametric test.

### In vivo administration of recombinant IL-30 exacerbates encephalitogenic inflammation

IL-30 can be secreted independently of Ebi3^32^. Therefore, aberrant susceptibility to EAE seen in the *Il27p28* gene deficient animals in this study may stem from defective IL-27 and/or IL-30 secretion from the targeted myeloid cells or microglia. We were particularly intrigued by the unexpected elevation of *Il27p28* mRNA expression in ‘wildtype’ CNS APC subsets within those cell type specific IL-30^-/-^ mice. As such increase did not occur in DC-specific IL-30^-/-^ mice where the disease severity remained unchanged, we posit that elevated *Il27p28* mRNA expression may be associated with severe disease progression. IL-30 may function as a natural antagonist of gp130-mediated signaling^32^. In support, it was previously reported that the secreted IL-30 could inhibit the biological functions of IL-27^22^. Therefore, IL-30 may antagonize anti-inflammatory roles of IL-27, driving encephalitogenic inflammation. To test this possibility, we induced EAE in B6 mice and then administered recombinant IL-27 or IL-30 via a mini-osmotic pump once the mice developed noticeable clinical signs. As shown in Fig 5a, IL-27 administered substantially dampened the EAE severity, as previously reported^33^. By contrast, IL-30 administration exacerbated the clinical severity of the recipient mice (Fig 5a). In support of the disease severity, CNS infiltrating CD4 T cells were diminished by IL-27 administration, while IL-30 administration substantially increased the infiltration (Fig 5b). Inflammatory cytokine expression by infiltrating CD4 T cells was similarly affected by IL-27 and IL-30 administered in vivo. TNFα-, IL-17-, GM-CSF-, and IFNγ-expressing CD4 T cell accumulation in the CNS was markedly diminished by IL-27, while the accumulation was dramatically increased following IL-30 administration (Fig 5c). Inflammatory chemokine mRNA expression in the CNS followed similar pattern and was significantly diminished by IL-27 but rather increased by IL-30 (Fig 5d). We previously showed that Foxp3^+^ Treg cells are the primary target cells of IL-27 in vivo and that IL-27 induces Lag3 expression in Treg cells^34^ Consistent with the previous findings, IL-27 administered increased Lag3 expression in Treg cells compared to that of sham treated controls, while IL-30 administration significantly diminished Treg cell expression of Lag3 (Fig 5e). Likewise, we observed that Treg cell Lag3 expression was significantly diminished in microglia-specific IL-30^-/-^ mice (Fig 5f), in which *Il27p28* mRNA expression was dysregulated and the mice developed severe EAE (Fig 4a and 4f). We measured IL-30 protein secretion from the CNS homogenates and found that the level was significantly increased in microglia-specific IL-30^-/-^ compared to that in wild type mice with EAE (Fig 5g). Unexpectedly, however, we also noticed elevated IL-27 heterodimer secretion from the CNS homogenates (Fig 5h). Since Treg cell Lag3 expression was lower in this condition (Fig 5f), these results suggest that IL-30 produced may compete with IL-27 for its binding to the receptor and to dampen inflammatory responses in part via Treg cells in vivo. The competition may result in elevated IL-27 levels in the CNS homogenates.

**Figure 5.**
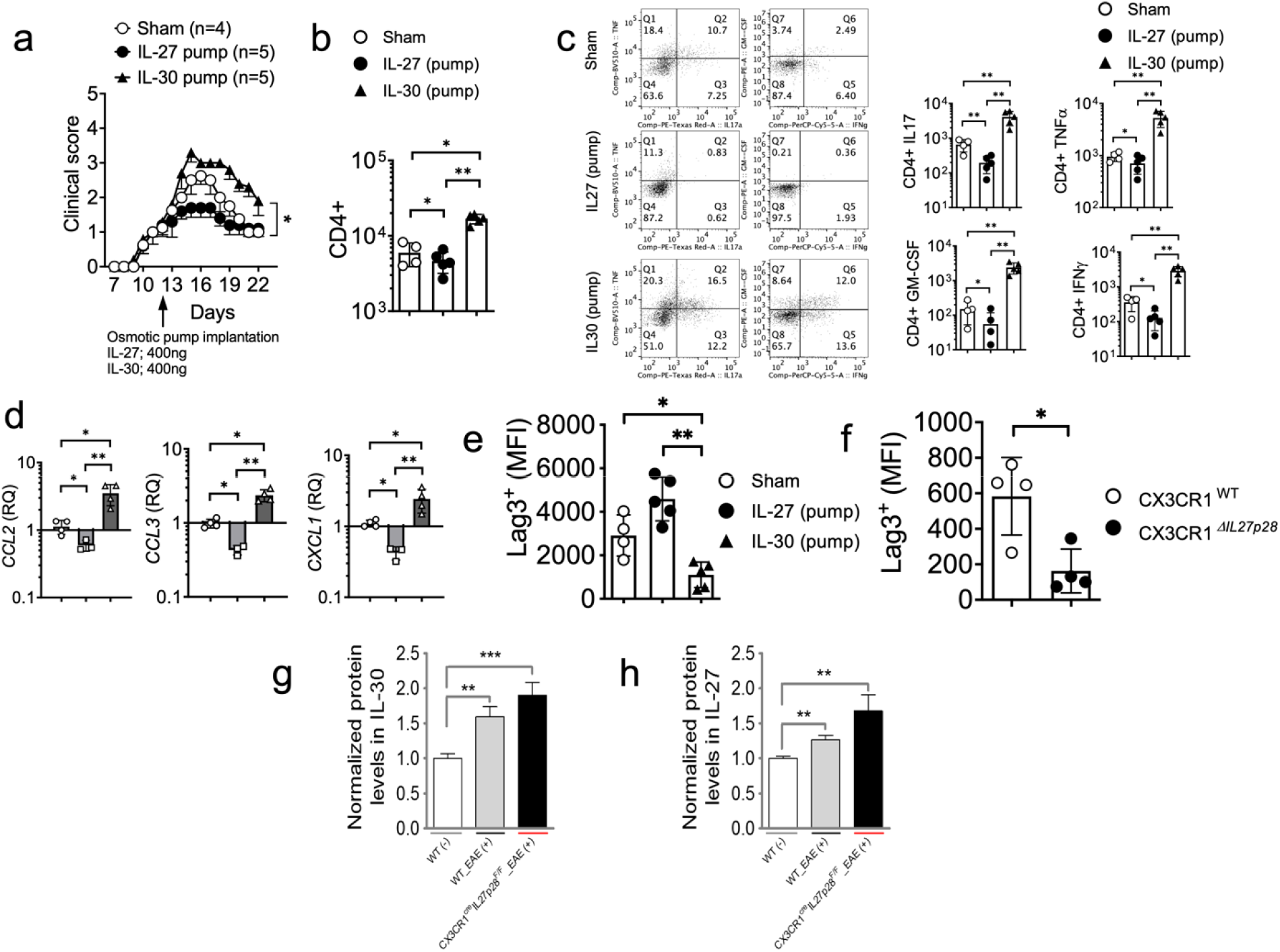
In vivo IL-30 administration develops significantly exacerbated EAE. C57BL/6 mice were induced for EAE. Osmotic pump containing IL-27 (400ng, n = 5), IL-30 (400ng, n = 5) were subcutaneously implanted or sham surgery (n = 4) was performed at 12 d post induction. (a) EAE score. (b) Total CD4^+^ T cell numbers in the CNS at 22 d post induction. (c) The numbers of GM-CSF, IFNγ, IL-17, and TNFα^+^ CD4 T cells were determined by intracellular cytokine staining at day 22 post immunization. (d) qPCR analysis of the indicated chemokine expression in the brain from sham, IL-27-pump and IL-30-pump group. Gene expression was normalized by *Gapdh* and compared to that of sham surgery group. (e) Lag3 expression of CNS infiltrating Treg cells was determined by flow cytometry. (f) Cx3cr1^WT^ and Cx3cr1^Δ*Il27p28*^ mice induced for EAE as described in Fig 4 were used to measure CNS infiltrating Treg cell expression of Lag3. (g and h) IL-30 and IL-27 levels in the CNS homogenates were measured by ELISA and were normalized to those of naïve wild type mice. N = 7 per group. *p < 0.05; **p < 0.01; ***p < 0.001; as determined by Mann-Whitney nonparametric test.

### IL-30 does not display regulatory properties in vitro

IL-30 fails to support T cell proliferation or IFNγ production^35^. However, IL-30 reportedly inhibits the production of IL-17 and IL-10 triggered by IL-27 or IL-6 stimulation in activated T cells^32,36^. In addition, IL-30 also induces LPS-induced TNFα and IP-10 production in monocytes^37^. Since we observed that IL-30 may be able to antagonize IL-27’s regulatory function in vivo, we sought to test if IL-30 expresses regulatory properties in vitro. Naïve CD4 and CD8 T cells were stimulated in the presence of recombinant IL-27 or IL-30. As expected, IL-27 rapidly phosphorylated both Stat1 and Stat3 in CD4 and CD8 T cells (Fig 6a and data not shown). By contrast, IL-30 stimulation had little effects on Stat phosphorylation (Fig 6a). The lack of Stat phosphorylation by IL-30 stimulation led us to further examine its ability to induce or to antagonize IFNγ or IL-10 expression in activated T cells. CD4 T cells stimulated under Th1 or Th17 polarization condition substantially upregulated *Ifng* mRNA expression in response to IL-27 but not to IL-30 (Fig 6b and data not shown). Likewise, IL-27 induced robust *Il10* mRNA expression in both developing Th1 and Th17 type CD4 T cells, while IL-30 failed to do so (Fig 6c and data not shown). Pre-stimulation with IL-27 effectively inhibited IL-27-induced Stat phosphorylation, whereas IL-30 pre-stimulation had no impact on interfering with IL-27-induced Stat phosphorylation (Fig 6d). Therefore, IL-30 alone does not appear to alter cytokine expression in activated T cells in vitro. IL-30 may influence T cell immunity through APCs, as *Ebi3* mRNA expression can be elevated especially when *Il27p28* mRNA expression was increased. In bone marrow-derived macrophages, neither IL-27 nor IL-30 had the ability to induce IL-12 or IL-27 expression (Supp Fig 5).

**Figure 6.**
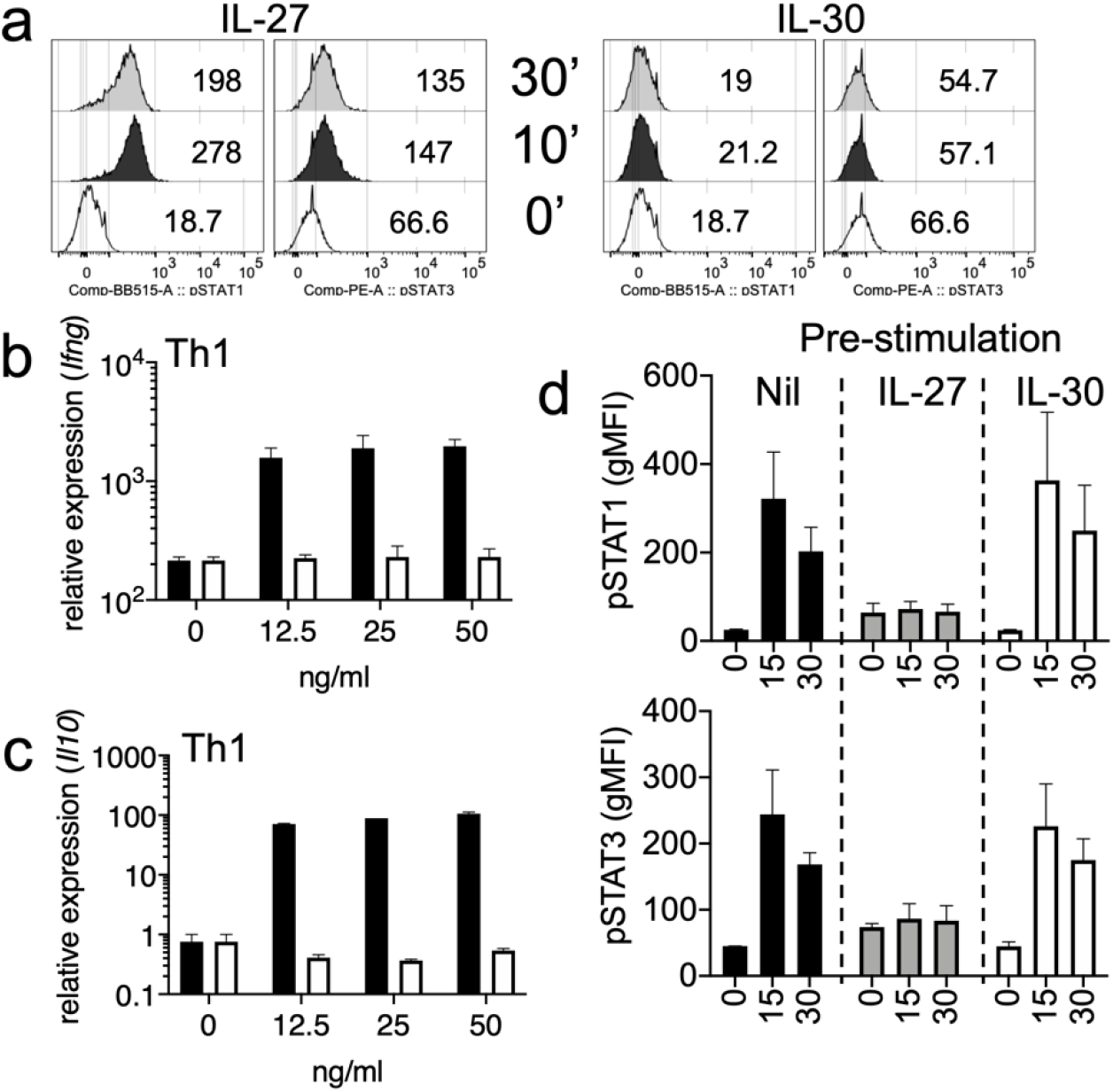
IL-30 stimulation in CD4 T cell activation in vitro. (a) FACS sorted CD4^+^ naïve cells were stimulated with recombinant IL-27, IL-30. Phosphorylated STAT1 and STAT3 expression was determined by flow cytometry at 10-and 30-minutes following stimulation. (b-c) Naive CD4 T cells were stimulated under Th1 polarization conditions in the presence of IL-27 or IL-30 (0-50 ng/ml) for 3 days. *Ifng and Il10* mRNA expression was determined by qPCR. (d) Naïve CD4 T cells were incubated with media (Nil), 50ng IL-27, or 50ng IL-30 for one hour. The cells were then washed and restimulated with IL-27. Stat1 and Stat3 phosphorylation was determined by flow cytometry at 15 and 30 minute following stimulation. The data shown are representative of two independent experiments.

## Discussion

In this study, we took advantage of cell type specific IL-30-deficient mouse models to identify the sources of IL-30 and to gain closer insights into the immune regulatory functions of IL-27 (and of IL-30 itself) in autoimmune inflammation. Consistent with the previous reports, we found that infiltrating myeloid cell-derived IL-30 is critical to limit autoimmune inflammation in the CNS. Interestingly, other CNS resident APC subsets, such as microglia, were equally important in modulating inflammatory responses within the CNS, as the lack of IL-30 expression in microglia also confers greater susceptibility to autoimmunity. Unexpectedly, we found that mice deficient in IL-30 specifically in DCs were not susceptible to the disease, suggesting a dispensable role of DC-derived IL-30 in EAE pathogenesis.

IL-30 was initially considered an IL-27-specific subunit; thus, mice deficient in or overexpressing the *Il27p28* gene have been used to interrogate the immune regulatory functions of IL-27. Mice overexpressing the *Il27p28* gene are resistant to autoimmune inflammation, and it was proposed that such resistance might originate from IL-27’s ability to antagonize inflammatory T cell responses, particularly Th1 and Th17 immunity^18^. Likewise, mice deficient in IL-30 are highly susceptible to EAE and express heightened Th17 immunity^5^. Therefore, it was naturally concluded that IL-30’s ability to control inflammatory responses is through its role as an IL-27 subunit. It was then discovered that IL-30 can be secreted in the absence of the Ebi3 subunit and is able to modulate immune responses^32^. Tagawa and colleagues reported that IL-30 alone can mediate immune regulatory functions, suppressing allogenic T cell responses^22^. More recently, Park et al. reported that IL-30 can act as a negative regulator of both B and T cell responses during *T. gondii* infection, independently of IL-27^30^. IL-30 may antagonize other cytokines, such as IL-27 or IL-6 that utilizes the gp130 surface receptor for the signaling^32^. Although IL-30 stimulation itself does not trigger any detectable Stat phosphorylation in vitro, its presence may be sufficient to hinder Stat1 and Stat3 phosphorylation induced by IL-27 or IL-6^32^. Consistent with these reports, we made a similar observation that IL-30 stimulation does not induce Stat phosphorylation in T cells. Unexpectedly, however, we also noticed that IL-30 fails to interfere with IL-27-induced Stat1/3 phosphorylation or IL-27-stimulated gene expression in vitro. Petes et al. previously reported that IL-30-induced Stat1 and Stat3 phosphorylation can be bimodal and delayed second phosphorylation event occurs at later time points in THP-1 cells^37^. When primary monocytes were used, delayed and weak Stat1 but not Stat3 phosphorylation was similarly observed^37^. However, we found no signs of delayed Stat phosphorylation in T cells (data not shown). Moreover, neither IL-27 nor IL-30 altered cytokine expression in bone marrow-derived macrophages. While the reason underlying the discrepancy is not clear, concentrations used may account for the difference, as the previous study used higher IL-30 concentration. IL-30 may be able to signal through gp130 homodimers without soluble IL-6Rα only at high concentrations^38^. The precise mechanisms by which IL-30 modulates inflammatory responses remain to be determined. Thus, IL-30 does not impact T cells and macrophages in vitro; however, our data suggest that IL-30 may act on other target cells, partly Treg cells, to modulate inflammatory responses in vivo.

Finding dysregulated *Il27p28* mRNA expression in some cell type specific IL-30^-/-^ mice with severe disease is of particular interest. What does trigger such an aberrant expression? IFNγ has previously been reported to induce IL-30 expression in myeloid cells^39^, and we likewise found elevated IFNγ expression in infiltrating CD4 T cells. IL-30 expression may thus be directly correlated with the disease severity and inflammation. Disproportionate expression of *Il27p28* mRNA may result in excessive IL-30, which may interfere with IL-27’s function to inhibit the inflammation. As a result, IFNγ production by autoreactive T cells continuously increases, further amplifying IL-30 expression. Although we found no evidence that IL-30 alone is able to induce Stat phosphorylation and gene expression in T cells in vitro, IL-30 may be able to do so in vivo, in part, based on its ability to antagonize IL-27’s function to modulate Treg cell Lag3 expression. Expression of Treg cell markers, ICOS, GITR, CD25, and Foxp3 remained unchanged in cell type-specific IL-30^-/-^ mice or in IL-27 or IL-30 treatment. We previously showed that Tim-3, CD39, and CTLA4 expression in Treg cells is not affected by the lack of IL-27 signaling^33^. However, we noticed that Lag3 expression on Treg cells may be balanced by the IL-27 and IL-30. Therefore, IL-30-induced effects may be operated in part by reducing Lag3 expression in Treg cells, although we cannot exclude the possibility that IL-30 itself poses unknown immune regulatory functions in vivo, because the receptor gp130 is expressed on multiple cell types including myeloid cells, B cells, and endothelial cells, etc^40^. Alternatively, IL-30 may form a complex with other subunits, such as cytokine-like factor 1 (CLF1) or soluble IL-6Rα^41^, and it is possible that different complexes may be preferentially formed depending on the type of cells. Whether these complexes mediate different functions remains to be determined. Of note, neutralizing or blocking antibodies against IL-30 cannot be used to investigate this possibility, because it will affect both IL-30 and IL-27. One may require a system in which IL-30 itself cannot be secreted unless it forms the heterodimeric IL-27 complex. Feige and colleagues recently reported a discrepant IL-30 secretion pattern between human and mice^42^. A single amino acid difference is sufficient to alter the secretion pattern of IL-30. Unlike mouse IL-30, which has two cysteine residues that enable a stable secretion of IL-30 independent of Ebi3, human IL-30 has only one cysteine residue. Thus, in the absence of Ebi3, IL-30 fails to be secreted; instead, it retains in the ER and ultimately gets degraded^42^, suggesting that Ebi3-independent IL-30 secretion will not occur in humans. We thus speculate that the in vivo functions of the IL-27:IL-30 axis are different in humans and mice. Further investigation will be necessary to examine the regulatory functions of IL-30.

DCs, especially XCR1^+^ cDC1 type subsets, express IL-30 when immunized with a combination adjuvant, poly I:C and agonistic anti-CD40 Ab^43^. Our finding that DC-derived IL-30 plays little role in limiting EAE pathogenesis and encephalitogenic immune responses suggests that DCs may not be the primary source of IL-30 during autoimmune inflammation in the CNS. DC-derived IL-30 may be important in the secondary lymphoid tissues during priming event as seen in the spleen following intravenous immunization with adjuvants^43^, and IL-30-derived from inflammatory monocytes/macrophages or tissue resident APC subsets may be more crucial in limiting immune responses in the target tissues.

In summary, we report that IL-30 expresses a regulatory property to promote autoimmune inflammation in vivo, which could partially be mediated by interfering with regulatory function of IL-27. Further investigation is needed to understand its precise immune modulatory functions, through which a novel approach targeting IL-30 and/or IL-27 may be developed.

## Materials and Methods

### Animals

C57BL/6, CD11c (*Itgax*)^Cre^ (strain #8068), *Gfap*^Cre^ (strain #24098), LysM (*Lyz2*)^Cre^ (strain #4781), and *Cx3cr1*^Cre^ (strain #25524) mice were purchased from the Jackson Laboratory (Bar Harbor, ME). *Il27p28*^fl/fl^ mice were previously reported^26^. All mice were bred in a specific pathogen-free facility at Northwestern University Feinberg School of Medicine. All the animal experiments were approved by the institutional animal care and use committees (IACUC) of Northwestern University (protocol #IS00015862).

### EAE induction

Mice were subcutaneously injected with 200 μL of an emulsion containing 300 μg of MOG_35-55_ peptide (BioSynthesis, Lewisville, TX) and equal volume of Complete Freund’s adjuvant supplemented with 5 mg/mL of Mycobacterium tuberculosis strain H37Ra (Difco, Detroit, MI). Additionally, mice were intraperitoneally injected with 200 ng of pertussis toxin (Sigma, St. Louis, MO) at the day of immunization and 48 h later. Disease development was analyzed daily and scored on a 0-5 scale: 0, no clinical signs; 1, limp tail, 2, hind limb weakness, 3, hind limb paralysis, 4, hind limb paralysis and front limb weakness, 5, moribund or death.

### Osmotic pump implantation

Mice anesthetized with Ketamine and Xylazine were subcutaneously implanted with a mini-osmotic pump (#1007D, Alzet Durect, Cupertino, CA) as previously described^11^. This pump system has a reservoir volume of 100 μl and allows for the continuous delivery of the content for 7 days without the need for external connections or frequent handling of animals. Pumps containing 400 ng of rIL-27 or rIL-30 (R&D Systems, Minneapolis, MN) were implanted at day 12 post immunization. Mice with sham surgery were used as controls.

### Flow Cytometry

Mononuclear cells from the CNS of EAE mice were isolated by Percoll gradient centrifugation as previously described^44^. The cells were then stained with anti-CD4 (RM4–5), anti-CD44 (IM7), anti-CD25 (PC61.5), anti-GITR (DTA-1), anti-Foxp3 (FJK-16s), and anti-ICOS (C398.4A) antibodies. For intracellular staining, harvested cells were stimulated ex vivo with PMA (10ng/mL, Millipore-Sigma) and ionomycin (1 μM, Millipore-Sigma) for 4 h in the presence of 2 μM monensin (Calbiochem) during the last 2 h of stimulation. Cells were immediately fixed with 4% paraformaldehyde, permeabilized, and stained with anti-IL-17 (TC11-18H10), anti-IFNγ (XMG1.2), anti-TNFα (TN3-19), anti-GM-CSF (MP1-22E9) antibodies. All the antibodies were purchased from eBioscience (San Diego, CA), BD PharMingen (San Diego, CA), and Biolegend (San Diego, CA). In some experiments cells were stimulated with recombinant IL-27 or IL-30 and phosphorylated Stat1 (4a) and Stat3 (LUVNKLA) expression was determined by flow cytometry. Samples were acquired using a FACSCelesta (BD Bioscience) and analyzed using a FlowJo (Treestar, Ashland, OR). Cytokine-expressing effector CD4 T cells were enumerated by flow cytometry. CNS APC subsets were sorted based on CD45 and CD11b expression using a FACSMelody cell sorter (BD Bioscience). Sorted cells were subjected to gene expression by qPCR as described below.

### Cytometric Beads Array

Serum cytokines were determined using Cytometric Beads Array (BD Biosciences) according to the manufacturer’s instructions. The data were analyzed using the CBA software, and the standard curve for each cytokine was generated using the mixed cytokine standard.

### Real-Time Quantitative PCR

Mice with EAE were euthanized and perfused with PBS. The brain and spinal cords were isolated and total RNA was extracted using a TRIzol reagent according to the manufacturer’s instructions (Invitrogen). In some experiments using bone marrow-derived macrophages, bone marrow cells harvested from the tibia and femur were cultured with L929-media (15% volume) for 6 days. Bone marrow generated macrophages were then harvested (>95% express CD11b). Harvested cells were then stimulated with IL-27 or IL-30, and cytokine gene expression was determined by qPCR as below. cDNA was then obtained using a MMLV reverse transcriptase (Promega, location). qPCR analysis was performed using a QuantStudio 3 Real-Time PCR System (Applied Biosystems, Waltham, MA) using a Radiant qPCR mastermix (Alkali Scientific, Fort Lauderdale, FL) or SYBR green mastermix (Applied Biosystems). The data were normalized by housekeeping *Gapdh* gene and then compared to the control group. Primers used for the study are listed in Supplementary Table 1.

### Sequential Protein extraction

The protein extraction was performed as previously reported^45^. In brief, mice were euthanized and perfused with cold PBS (10 mM, pH 7.4). The brains and spinal cords were then removed. Soluble protein fractions were obtained from the whole cortex and spinal cord using sequential protein extraction. Fractions were obtained by homogenization of the cortex and spinal cord with a dounce homogenizer in buffer including protease inhibitor cocktail (2 mL/200 μg of tissues; Thermo Scientific, 1862209). After centrifugation for 1.5h at 43.000 rpm in tube (#349622, 3.5 mL, 13 x 51mm tubes) for high-speed centrifugation from Beckman-Coulter, the supernatants were obtained and aliquoted and stored at −80 °C. To measure cytokines in the brain and spinal cord soluble fractions, IL-30 (R&D Systems, Minneapolis, MN) and IL-27 (LEGEND MAXTM, BioLegend) ELISA kits were used according to the manufacturer’s instructions.

### Statistical Analysis

Statistical significance was determined by the Mann-Whitney test using Prism software (GraphPad, San Diego, CA). p<0.05 was considered statistically significant.

## Author Contributions

D.K. designed and performed most of the experiments, analyzed the data, and wrote the manuscript. S.K. and M.K. performed experiments shown in part of the Figures 5 and 6. Z.Y. provided key reagents. B.M. designed the experiments, analyzed the data, and wrote the manuscript. All authors reviewed the manuscript.

## Bibliography

1 Kourko, O., Seaver, K., Odoardi, N., Basta, S. & Gee, K. IL-27, IL-30, and IL-35: A Cytokine Triumvirate in Cancer. Front Oncol 9, 969, doi:10.3389/fonc.2019.00969 (2019).

2 Yoshida, H. & Hunter, C. A. The immunobiology of interleukin-27. Annu Rev Immunol 33, 417–443, doi:10.1146/annurev-immunol-032414-112134 (2015).

3 Yoshimura, T. et al. Two-sided roles of IL-27: induction of Th1 differentiation on naive CD4+ T cells versus suppression of proinflammatory cytokine production including IL-23-induced IL-17 on activated CD4+ T cells partially through STAT3-dependent mechanism. J Immunol 177, 5377–5385, doi:10.4049/jimmunol.177.8.5377 (2006).

4 Cao, Y., Doodes, P. D., Glant, T. T. & Finnegan, A. IL-27 induces a Th1 immune response and susceptibility to experimental arthritis. J Immunol 180, 922–930, doi:10.4049/jimmunol.180.2.922 (2008).

5 Diveu, C. et al. IL-27 blocks RORc expression to inhibit lineage commitment of Th17 cells. J Immunol 182, 5748–5756, doi:10.4049/jimmunol.0801162 (2009).

6 Pot, C. et al. Cutting edge: IL-27 induces the transcription factor c-Maf, cytokine IL-21, and the costimulatory receptor ICOS that coordinately act together to promote differentiation of IL-10-producing Tr1 cells. J Immunol 183, 797–801, doi:10.4049/jimmunol.0901233 (2009).

7 Zhang, H. et al. An IL-27-Driven Transcriptional Network Identifies Regulators of IL-10 Expression across T Helper Cell Subsets. Cell Rep 33, 108433, doi:10.1016/j.celrep.2020.108433 (2020).

8 Pot, C., Apetoh, L. & Kuchroo, V. K. Type 1 regulatory T cells (Tr1) in autoimmunity. Semin Immunol 23, 202–208, doi:10.1016/j.smim.2011.07.005 (2011).

9 Zeng, H., Zhang, R., Jin, B. & Chen, L. Type 1 regulatory T cells: a new mechanism of peripheral immune tolerance. Cell Mol Immunol 12, 566–571, doi:10.1038/cmi.2015.44 (2015).

10 Batten, M. et al. Interleukin 27 limits autoimmune encephalomyelitis by suppressing the development of interleukin 17-producing T cells. Nat Immunol 7, 929–936, doi:10.1038/ni1375 (2006).

11 Kim, D. et al. Cutting Edge: IL-27 Attenuates Autoimmune Neuroinflammation via Regulatory T Cell/Lag3-Dependent but IL-10-Independent Mechanisms In Vivo. J Immunol 202, 1680–1685, doi:10.4049/jimmunol.1800898 (2019).

12 Molle, C. et al. IL-27 synthesis induced by TLR ligation critically depends on IFN regulatory factor 3. J Immunol 178, 7607–7615, doi:10.4049/jimmunol.178.12.7607 (2007).

13 Blahoianu, M. A., Rahimi, A. A., Kozlowski, M., Angel, J. B. & Kumar, A. IFN-gamma-induced IL-27 and IL-27p28 expression are differentially regulated through JNK MAPK and PI3K pathways independent of Jak/STAT in human monocytic cells. Immunobiology 219, 1–8, doi:10.1016/j.imbio.2013.06.001 (2014).

14 Rajaiah, R., Puttabyatappa, M., Polumuri, S. K. & Moudgil, K. D. Interleukin-27 and interferongamma are involved in regulation of autoimmune arthritis. J Biol Chem 286, 2817–2825, doi:10.1074/jbc.M110.187013 (2011).

15 Ramgolam, V. S., Sha, Y., Jin, J., Zhang, X. & Markovic-Plese, S. IFN-beta inhibits human Th17 cell differentiation. J Immunol 183, 5418–5427, doi:10.4049/jimmunol.0803227 (2009).

16 Di Carlo, E. Decoding the Role of Interleukin-30 in the Crosstalk Between Cancer and Myeloid Cells. Cells 9, doi:10.3390/cells9030615 (2020).

17 Min, B., Kim, D. & Feige, M. J. IL-30(dagger) (IL-27A): a familiar stranger in immunity, inflammation, and cancer. Exp Mol Med 53, 823–834, doi:10.1038/s12276-021-00630-x (2021).

18 Chong, W. P. et al. IL-27p28 inhibits central nervous system autoimmunity by concurrently antagonizing Th1 and Th17 responses. J Autoimmun 50, 12–22, doi:10.1016/j.jaut.2013.08.003 (2014).

19 Park, J. et al. Impact of Interleukin-27p28 on T and B Cell Responses during Toxoplasmosis. Infect Immun 87, doi:10.1128/IAI.00455-19 (2019).

20 Di Meo, S. et al. Interleukin-30 expression in prostate cancer and its draining lymph nodes correlates with advanced grade and stage. Clin Cancer Res 20, 585–594, doi:10.1158/1078-0432.CCR-13-2240 (2014).

21 Sorrentino, C. et al. Interleukin-30/IL27p28 Shapes Prostate Cancer Stem-like Cell Behavior and Is Critical for Tumor Onset and Metastasization. Cancer Res 78, 2654–2668, doi:10.1158/0008-5472.CAN-17-3117 (2018).

22 Shimozato, O. et al. The secreted form of p28 subunit of interleukin (IL)-27 inhibits biological functions of IL-27 and suppresses anti-allogeneic immune responses. Immunology 128, e816–825, doi:10.1111/j.1365-2567.2009.03088.x (2009).

23 Korn, T. et al. Myelin-specific regulatory T cells accumulate in the CNS but fail to control autoimmune inflammation. Nat Med 13, 423–431, doi:10.1038/nm1564 (2007).

24 Dibra, D., Cutrera, J. J. & Li, S. Coordination between TLR9 signaling in macrophages and CD3 signaling in T cells induces robust expression of IL-30. J Immunol 188, 3709–3715, doi:10.4049/jimmunol.1100883 (2012).

25 Awasthi, A. et al. A dominant function for interleukin 27 in generating interleukin 10-producing anti-inflammatory T cells. Nat Immunol 8, 1380–1389, doi:10.1038/ni1541 (2007).

26 Zhang, S. et al. High susceptibility to liver injury in IL-27 p28 conditional knockout mice involves intrinsic interferon-gamma dysregulation of CD4+ T cells. Hepatology 57, 1620–1631, doi:10.1002/hep.26166 (2013).

27 Wei, J. et al. Critical role of dendritic cell-derived IL-27 in antitumor immunity through regulating the recruitment and activation of NK and NKT cells. J Immunol 191, 500–508, doi:10.4049/jimmunol.1300328 (2013).

28 de Aquino, M. T. et al. IL-27 limits central nervous system viral clearance by promoting IL-10 and enhances demyelination. J Immunol 193, 285–294, doi:10.4049/jimmunol.1400058 (2014).

29 Sonobe, Y. et al. Production of IL-27 and other IL-12 family cytokines by microglia and their subpopulations. Brain Res 1040, 202–207, doi:10.1016/j.brainres.2005.01.100 (2005).

30 Prinz, M. et al. Distinct and nonredundant in vivo functions of IFNAR on myeloid cells limit autoimmunity in the central nervous system. Immunity 28, 675–686, doi:10.1016/j.immuni.2008.03.011 (2008).

31 Yona, S. et al. Fate mapping reveals origins and dynamics of monocytes and tissue macrophages under homeostasis. Immunity 38, 79–91, doi:10.1016/j.immuni.2012.12.001 (2013).

32 Stumhofer, J. S. et al. A role for IL-27p28 as an antagonist of gp130-mediated signaling. Nat Immunol 11, 1119–1126, doi:10.1038/ni.1957 (2010).

33 Do, J. et al. Treg-specific IL-27Ralpha deletion uncovers a key role for IL-27 in Treg function to control autoimmunity. Proc Natl Acad Sci U S A 114, 10190–10195, doi:10.1073/pnas.1703100114 (2017).

34 Do, J. S. et al. An IL-27/Lag3 axis enhances Foxp3+ regulatory T cell-suppressive function and therapeutic efficacy. Mucosal Immunol 9, 137–145, doi:10.1038/mi.2015.45 (2016).

35 Pflanz, S. et al. IL-27, a heterodimeric cytokine composed of EBI3 and p28 protein, induces proliferation of naive CD4+ T cells. Immunity 16, 779–790, doi:10.1016/s1074-7613(02)00324-2 (2002).

36 Stumhofer, J. S. et al. Interleukin 27 negatively regulates the development of interleukin 17-producing T helper cells during chronic inflammation of the central nervous system. Nat Immunol 7, 937–945, doi:10.1038/ni1376 (2006).

37 Petes, C., Mariani, M. K., Yang, Y., Grandvaux, N. & Gee, K. Interleukin (IL)-6 Inhibits IL-27-and IL-30-Mediated Inflammatory Responses in Human Monocytes. Front Immunol 9, 256, doi:10.3389/fimmu.2018.00256 (2018).

38 Garbers, C. et al. An interleukin-6 receptor-dependent molecular switch mediates signal transduction of the IL-27 cytokine subunit p28 (IL-30) via a gp130 protein receptor homodimer. J Biol Chem 288, 4346–4354, doi:10.1074/jbc.M112.432955 (2013).

39 Murugaiyan, G., Mittal, A. & Weiner, H. L. Identification of an IL-27/osteopontin axis in dendritic cells and its modulation by IFN-gamma limits IL-17-mediated autoimmune inflammation. Proc Natl Acad Sci US A 107, 11495–11500, doi:10.1073/pnas.1002099107 (2010).

40 Tanaka, T. et al. Interleukin-27 induces the endothelial differentiation in Sca-1+ cardiac resident stem cells. Cytokine 75, 365–372, doi:10.1016/j.cyto.2015.06.009 (2015).

41 Crabe, S. et al. The IL-27 p28 subunit binds cytokine-like factor 1 to form a cytokine regulating NK and T cell activities requiring IL-6R for signaling. J Immunol 183, 7692–7702, doi:10.4049/jimmunol.0901464 (2009).

42 Muller, S. I. et al. A folding switch regulates interleukin 27 biogenesis and secretion of its alpha-subunit as a cytokine. Proc Natl Acad Sci US A 116, 1585–1590, doi:10.1073/pnas.1816698116 (2019).

43 Kilgore, A. M. et al. IL-27p28 Production by XCR1(+) Dendritic Cells and Monocytes Effectively Predicts Adjuvant-Elicited CD8(+) T Cell Responses. Immunohorizons 2, 1–11, doi:10.4049/immunohorizons.1700054 (2018).

44 Hwang, M. et al. Distinct CD4 T-cell effects on primary versus recall CD8 T-cell responses during viral encephalomyelitis. Immunology 144, 374–386, doi:10.1111/imm.12378 (2015).

45 Boza-Serrano, A., Yang, Y., Paulus, A. & Deierborg, T. Innate immune alterations are elicited in microglial cells before plaque deposition in the Alzheimer’s disease mouse model 5xFAD. Sci Rep 8, 1550, doi:10.1038/s41598-018-19699-y (2018).

