## Supplementary Figures and Table for "Cell type specific IL-27p28 (IL-30) deletion uncovers an unexpected regulatory function of IL-30 in autoimmune inflammation"

Supp Figure 1

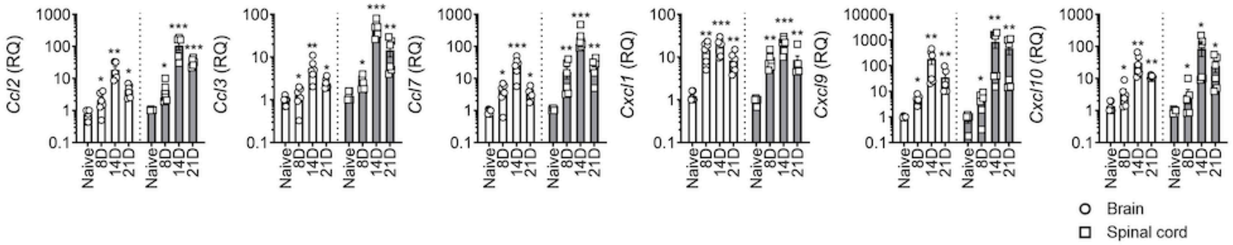


**Supplementary Figure 1. Expression of chemokines mRNA in EAE**.
Expression of the *Ccl2, Ccl3, Ccl7, Cxcl1, Cxcl9* and *Cxcl10* mRNA in the brain and spinal cords were determined at the time points shown in Figure 1. Data normalized to *Gapdh* gene were compared to those of naive mice. n = 4-6 per group. *p < 0.05; **p < 0.01; ***p < 0.001; ****p<0.0001; as determined by Mann-Whitney nonparametric test.

Supp Figure 2

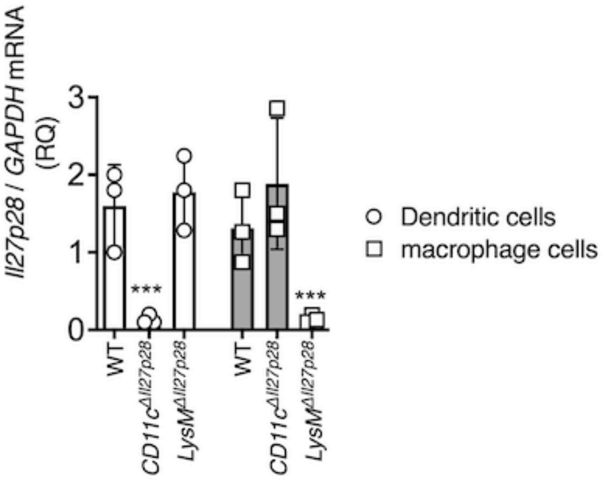


**Supplementary Figure 2. IL27p28 deletion in CD11c$^{\Delta Il27p28}$ *and* LysM$^{\Delta Il27p28}$ *mice*.** FACS sorted CD11c$^{+}$ (dendritic cells), CD11b$^{+}$CD45$^{+}$ (macrophages) cells from spleen and analyzed for the expression of *Il27p28* by qPCR. Data were normalized to WT mice. n = 3 per group. *$p < 0.05$; **$p < 0.01$; ***$p < 0.001$; as determined by Mann-Whitney nonparametric test.

Supp Figure 3

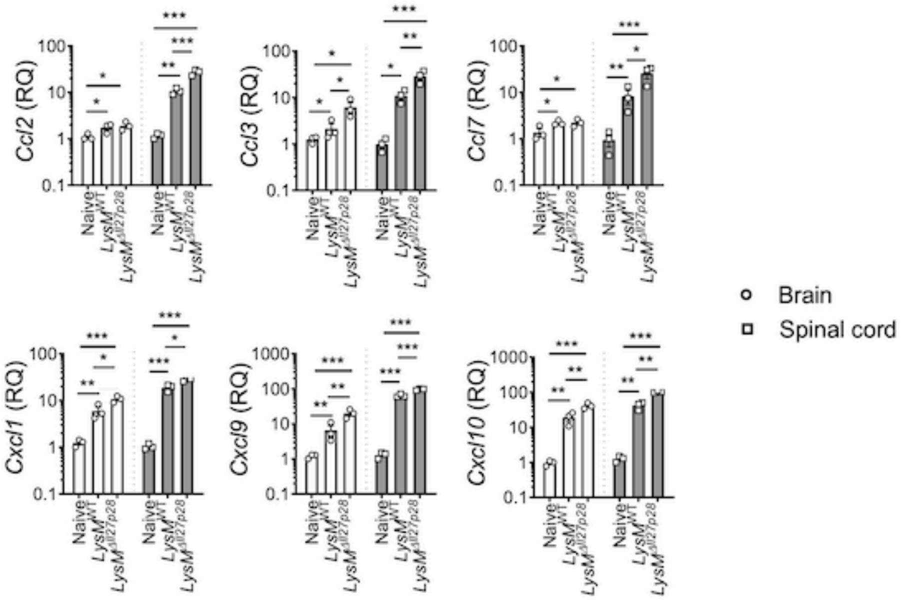


**Supplementary Figure 3. Chemokine expression in LysM$^{\Delta Il27p28}$ mice with EAE.** Gene expression was determined in brain and spinal cords extracts by qPCR 17 days after immunization. Gene expression was normalized to *Gapdh* gene. n = 3 per group. *p < 0.05; **p < 0.01; ***p < 0.001; as determined by Mann-Whitney nonparametric test.

Supp Figure 4

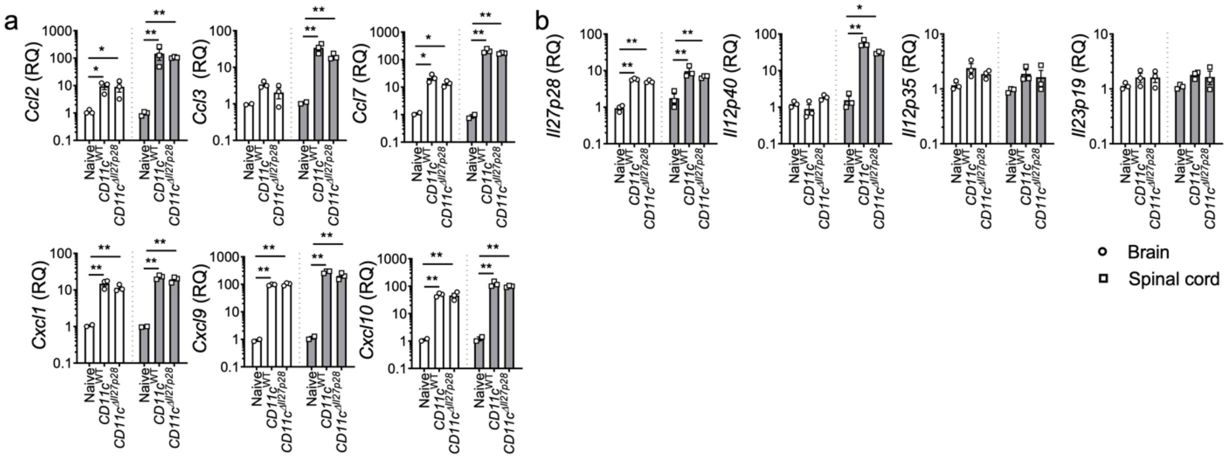


**Supplementary Figure 4. Chemokine and cytokine genes expression in CD11c$^{\Delta Il27p28}$ mice**. Gene expression was determined from the brain and spinal cord extracts by qPCR 17 days after immunization. Data normalized to *Gapdh* gene were compared to those of naive mice. n = 3 per group. *p < 0.05; **p < 0.01; ***p < 0.001; as determined by Mann-Whitney nonparametric test.

Supp Figure 5

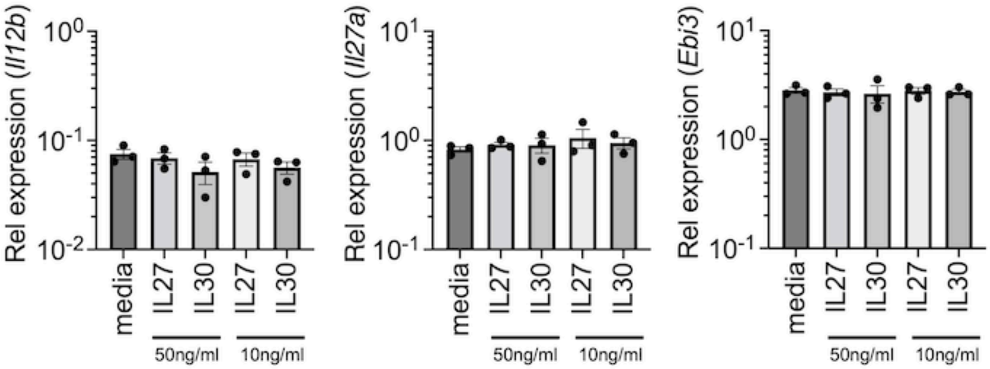


**Supplementary Figure 5. IL-27 and IL-30 stimulation of bone marrow derived macrophages.** Bone marrow derived macrophages generated from B6 mice were stimulated with recombinant IL-27 or IL-30 (10ng/ml or 50ng/ml) for 4 hours. Indicated gene expression was determined by qPCR. The data shown are the mean and SEM of three independent experiments.

Supp Table 1

| Designed Primers | |
|---|---|
| **Ccl2** | |
| For | 5' GTTAACGCCCCACTCACCT 3' |
| Rev | 5' AAAAACTACAGCTTCTTTGGGACACCT 3' |
| **Ccl3** | |
| For | 5' GTACCATGACACTCTGCAACC3' |
| Rev | 5' GTCAGGAAAATGACACCTGGCTG3' |
| **Ccl7** | |
| For | 5' TGAAAACCCCAACTCCAAAG3' |
| Rev | 5' CATTCCTTAGGCGTGACCAT3' |
| **Cxcl1** | |
| For | 5' CCGAAGTCATAGCCACACTCAA3' |
| Rev | 5' GCAGTCTGTCTTCTTTCTCCGTTA3' |
| **Cxcl9** | |
| For | 5' AGAACGGTGCGCTGCAC3' |
| Rev | 5' CCTATGGCCCTGGGTCTCA3' |
| **Cxcl10** | |
| For | 5' GCCGTCATTTTCTGCCTCAT3' |
| Rev | 5' GCTTCCCTATGGCCCTCATT3' |
| **Gapdh** | |
| For | 5' ATGCCTGCTTCACCACCTTCT3' |
| Rev | 5' CATGGCCTTCCGTGTTCCTA3' |
